# Individual variation in the avian gut microbiota: the influence of host state and environmental heterogeneity

**DOI:** 10.1101/2022.09.05.506623

**Authors:** Shane E. Somers, Gabrielle L. Davidson, Crystal N. Johnson, Michael S. Reichert, Jodie M. S. Crane, R. Paul Ross, Catherine Stanton, John L. Quinn

## Abstract

The gut microbiome has important consequences for fitness, yet the complex, interactive nature of ecological factors that influence the gut microbiome has scarcely been investigated in natural populations. We sampled the gut microbiota of wild great tits (Parus major) at different life stages and across multiple conifer and mixed woodland fragments, allowing us to evaluate multiple factors that relate to within-individual gut microbiota acquisition, including habitat type, nest position and life history traits. The gut microbiota varied with both environment and life-history in ways that were largely dependent on age. Notably, it was the nestling, as opposed to the adult gut microbiota that was most sensitive to ecological variation, pointing to a high degree of developmental plasticity. Individual nestling differences in gut microbiota were consistently different (repeatable) from one to two weeks of life, driven entirely by the effect of sharing the same nest. Our findings point to important early developmental windows in which the gut microbiota are most sensitive to environmental variation and suggest reproductive timing, and hence parental quality or food availability, interact with the microbiome.

## Introduction

The gut microbiome―the enteric microbial community and their genes―is strongly influenced by the environment (Rothschild et al., 2018), and plays a major role in host ecology and evolution (Dethlefsen et al., 2007; Koskella et al., 2017). Although much progress has been made in understanding the interplay between gut microbiome communities and host phenotypes, this research has been primarily restricted to lab systems (Colston & Jackson, 2016; G. L. Davidson et al., 2020). In nature, animal microbiomes are far more variable and diverse than their captive counterparts because they are exposed to a much wider variety of conditions (Hird, 2017; McKenzie et al., 2017), and because wild hosts are more genetically, and therefore biologically diverse than lab-reared animals. Exploring how gut microbiomes are acquired, their stability across life stages, and their interactions with extrinsic and intrinsic traits in nature are important for understanding what drives host phenotypic consequences associated with gut microbiome variation.

The gut microbiota can directly affect the development of key physiological and biological traits of the host (Cox et al., 2014; J. A. Foster et al., 2017; Hansen et al., 2012). Although most of our understanding of these processes comes from germ-free lab models and gut microbiota interventions, evidence is emerging that the developing gut microbiome has important consequences for hosts in natural systems. We previously showed in a wild-bird population that nestling gut microbiota diversity predicts nestling weight (G. L. Davidson et al., 2021), itself a key predictor of offspring survival and recruitment. Disruption to microbial gut colonisation at critical developmental periods can affect host immunity, metabolism and physical development in other systems such as bees, birds and frogs (Knutie et al., 2017; Schwarz et al., 2016; Simon et al., 2016; Warne et al., 2019). Early life is a critical time for gut colonisation and the developing gut microbiota in young animals is generally more sensitive to environmental variation than in the more established adult gut microbiota (Derrien et al., 2019; Grond et al., 2017). Although it is well known that the gut microbiota changes with development, the role that ecology plays in this process remains poorly understood.

Many fundamental drivers of ecological variation have the potential to affect the gut microbiota. Individual variation in the gut microbiota may be influenced by exposure to different environmental pools of microbes, for example, those in the air, soil or diet (Grond et al., 2017; Kartzinel et al., 2019; Liddicoat et al., 2020; Ren et al., 2017). The link with diet, in particular, is likely to explain why gut microbiota can vary with habitat type (Drobniak et al., 2021; Goossens et al., 2022; Teyssier, Rouffaer, et al., 2018). Vegetation and diet can also vary dramatically within habitats, for example with respect to edge effects (J. Chen et al., 1992; Wilkin et al., 2007), although it is unknown whether individual microbial communities are affected by these fine-scale processes. Between-seasonal dietary change has been linked to temporal variation in the gut microbiota (Baniel et al., 2021; Hicks et al., 2018; Maurice et al., 2015; Ren et al., 2017). Within-season variation in the gut microbiota has yet to be documented but is likely important because in many animals, reproductive timing is critical to ensure breeding coincides with seasonal peaks in food abundance (Brommer et al., 2002; Grüebler & Naef-Daenzer, 2010; Rodríguez et al., 2016; Rubenstein & Wikelski, 2003). Hence, the timing of reproduction should dictate nestlings’ exposure to dietary microbes, although this has never been tested in the wild.

Similarly indirect effects mediated by the host, for example, homeostatic responses to environmental stressors may have knock-on effects on the gut microbiota (Noguera et al., 2018; Stothart et al., 2019). Exercise intensity (Mailing et al., 2019) and energetics (reviewed in Lindsay et al. (2020)) are associated with the microbiota, while stress is also directly linked to the microbiota in general (Sudo et al., 2004), and in the context of reproduction specifically (MacLeod et al., 2022). Similarly, the gut microbiota in adults is mediated by a variety of sex hormones (Mallott et al., 2020), and can lead to sex-specific associations between the gut microbiota and environmental sources of variation (Org et al., 2016). For nestlings, larger brood sizes lead to increased competition for food (Smith et al., 1989), and higher glucocorticoid hormone concentrations (Greggor et al., 2017; Smith et al., 1988), both of which have been reported to influence their gut microbiota in the wild (David et al., 2014; G. Davidson et al., 2020; Noguera et al., 2018; Teyssier et al., 2020). At the same time, brood size is an indicator of parental effort (Pettifor et al., 2001), and alters glucocorticoid concentrations in adults (Bonier et al., 2011), which likewise could affect the adult microbiota.

Thus, the composition of gut microbes are expected to vary across time, in response to environmental variation, and with respect to a whole variety of individual state variables. Nevertheless, at the same time, individuality and stability in microbiota can also be important, as observed in studies of human health (L. Chen et al., 2021; Fassarella et al., 2021). Consistent among-individual variation indicates limited individual plasticity, that is, major constraints caused by intrinsic differences among individuals in some causal factor―for example, genetic, epigenetic, or early environment effects (Rajilić-Stojanović et al., 2013; Zoetendal et al., 2001). Typically stability in phenotypic variation among individuals is quantified by taking multiple measures from the same individual over time, and estimating individual repeatability using individual as a random effect in a mixed model framework (Nakagawa & Schielzeth, 2010). Mixed models also enable an examination of whether individual differences are independent of other individual or environmental factors that can also be included as random or fixed effects in the same model. Despite its widespread use in animal personality and cognition literature (Bell et al., 2009; Cauchoix et al., 2018), to our knowledge this approach has yet to be applied to identify the sources of variation in the gut microbiota.

We explored how the gut microbiota varied with the environment, with development, and with intrinsic factors using a generalist passerine bird species breeding in nest boxes across heterogenous and fragmented woodlands. The great tit (*Parus major*) is a widely used model organism in ecological and behavioural studies (Cole & Quinn, 2012; Drent et al., 2003; O’Shea et al., 2018; Tinbergen & Boerlijst, 1990). Within a single breeding season, we simultaneously investigated how the gut microbiota were related to host state (age, sex, reproductive effort) and a range of environmental factors (habitat type, distance to forest edge, number of siblings in the nest), and whether these explained consistent between-individual differences in the gut microbiota. While we had clear *a priori* reasons to expect environmental and life history traits to covary with gut microbiota variation, and to do so differentially across life stages because the developing gut microbiota in young animals are generally more sensitive to environmental variation than in the more established adult gut microbiota (Derrien et al., 2019; Grond et al., 2017), we had no *a priori* predictions about the direction of such effects in the context of gut microbiota community metrics (i.e. high vs low diversity and relative taxonomic abundance), owing to the complexity of host-microbe interactions (Douglas, 2018; K. R. Foster et al., 2017; Zaneveld et al., 2017). Finally, we conducted a range of repeatability analyses at the individual, the nest, and the woodland levels to determine whether individual nestlings differed consistently over a short but critical period of their development, from eight to 15 days old. We further examined whether any consistency at the individual-level was driven by intrinsic differences among individuals, or instead by shared nest or woodland effects, or indeed a range of other fixed effects as discussed above. Our analyses suggest a diverse and complex relationship between individual gut microbiota and the environment, the implications of which we suggest may well be substantial for populations.

## Methods

### Field monitoring and microbiota sampling

Birds were sampled from nine small woodland fragments across Co. Cork, Ireland, five of which were mixed/deciduous and four coniferous woodlands (see O’Shea et al. (2018). We collected 262 faecal samples from 204 great tits from 63 nests (see table S1 for final sample sizes post bioinformatic processing) for 16S rRNA gene sequencing. Nest boxes were monitored during April-June 2016 to determine lay dates, hatching dates and nestling survival. Typically, all individuals in a nest were sampled. Individual nestlings were sampled when they were 8 days old (+/− 1 day), and again, if they survived, when they were 15 days old (D8 and D15 birds respectively), at which point parents were also sampled. Birds were placed individually into sterile holding bags inside a heated holding case and naturally-produced faecal samples were collected. Urea has the potential to affect downstream sequencing and was removed through absorption by coffee filters placed as lining in the sampling bags (Khan et al., 1991). The faecal matter was collected within 15-20 minutes of placing birds in the sampling bag, after which birds were returned to the nest. Faecal sacks were ruptured immediately using a sterile inoculation loop and placed in a microcentrifuge tube containing 500uL of 100% ethanol. Samples were stored at −20°C within 8 hours of collection until DNA extraction. D8 nestlings were weighed, and individually identified by clipping the tip of one of their nails, taking care to avoid the blood vessel. Nestling birds were again weighed at D15 and ringed with a unique identifiable metal ring (British Trust for Ornithology). Samples from nestlings for which we had multiple samples (i.e. for both D8 and Day-15) are referred to as ‘repeat samples’. Adult birds were trapped on the nest and, if not already ringed, were fitted with a British Trust for Ornithology ring, weighed, and aged as either ‘immature’ (first year breeding) or ‘mature’ (second year/+ breeding) using plumage indicators.

### DNA Extractions

DNA was extracted from the dried faecal contents of all birds using the Qiagen QIAamp DNA Stool Kit, following the ‘Isolation of DNA from Stool for Pathogen Detection’ protocol (June 2012 edition), with modifications described in Shutt et al. (2020) to accommodate dried avian faeces. DNA was stored at −20°C. Full extraction methods are described in the Supporting Information of Davidson et al. (2021).

### Illumina MiSeq sequencing

Full library preparation details are described in Supporting Information of Davidson et al. (2021). Briefly, the V3-V4 variable region of the 16S rRNA gene was amplified from the DNA extracts using the 16S metagenomic sequencing library protocol (Illumina). The DNA was amplified with primers specific to the V3-V4 region of the 16S rRNA gene which also incorporates the Illumina overhang adaptor. Samples were sequenced on the MiSeq sequencing platform (Clinical Microbiomics, Denmark), using a 2 × 300 cycle kit, following standard Illumina sequencing protocols.

### Bioinformatics

The DADA2 pipeline was used to process the raw sequencing data (Callahan et al., 2016) in R version 3.5 (R Core Team, 2019). Sequence quality was visually inspected. Sequences were trimmed to remove adapters and lower quality reads (median quality scores below 25-30 threshold) at the extremities of the sequence and filtered to remove sequences with expected errors above 1. Read errors were estimated before dereplication. Forward and reverse reads were merged to construct ‘contig’ sequences, which were used to construct a sequence table of Amplicon Sequence Variants (ASV’s), which in turn counts the number of times each unique sequence is detected. The previous steps were performed for each run separately. Then the separate sequence tables were merged and chimeras removed using the ‘consensus’ method. Taxonomy was assigned to each ASV by RDP’s Naive Bayes Classifier (Wang et al., 2007) against the Silva reference database (version 132) (Quast et al., 2012). This method groups sequences with 100% sequence identity in contrast to the lower resolution OTU method which groups sequences at 97% identity. ASV’s allow greater sensitivity and specificity, better discrimination of ecological patterns than OTU’s and are reusable across studies (Callahan et al., 2017).

The DADA2 outputs were assembled into a single Phyloseq object (McMurdie & Holmes, 2013). Sequences identified as mitochondrial or chloroplast were removed. Sample completeness curves were plotted using vegan (Oksanen et al., 2019) and helped determine the lower cut-off for sample reads at 10,000 reads. Low read samples (<10,000 reads, 11 samples) were removed leaving 195 (adult=51, Day-8=81, Day-15=114) samples for the analysis. Alpha diversity (both Shannon and Chao1 diversity) was calculated using the ‘*estimate_richness’* function from the phyloseq package on the filtered dataset.

After removing low read samples (<10,000 reads, n = 11), chloroplast sequences and mitochondria sequences, there were 18,890,006 total reads clustered into 54,343 ASV’s in 246 samples (see table S1 for sample breakdown). Reads ranged from 10,220 to 557,336, with a mean of 76,789 reads per sample.

#### Statistical analysis

##### Datasets

The analyses detailed below were conducted on one of two subsets of the data, depending on the questions being explored. The first subset, referred to as the ‘*all birds*’ subset, contained birds of all developmental ages (8 day-old nestlings (D8); 15 day-old nestlings (D15); adults), and was used to examine the relationship between the gut microbiome, developmental age, and the interaction between developmental age and other life-history and environmental variables. Because it is generally expected that young animals are especially sensitive to environmental variation (Derrien et al., 2019; Grond et al., 2017), our main focus was on the interactions between developmental age and the other main effects. However we also explored the possibility that all main effects might have had interacting effects on microbiota. Analyses using the *all birds* dataset did not distinguish between birds breeding in their first year of life (immature) and those that were older (mature) because of convergence issues. The second subset of data contained adults only, and focussed on sex and its interaction with other life-history and environmental variables. All analyses were conducted in R version 3.6.3 (R Core Team, 2020).

##### Alpha diversity

Linear mixed models (LMM) were used to test the effect of host and environmental factors on alpha diversity. Shannon diversity (Shannon, 1948) measured richness weighted by abundance (the evenness of a community), and Chao1 (Chao, 1984) measured richness, specifically estimating taxa abundance and rare taxa missed from under sampling. Models were fit using the *lme4* package (Bates et al., 2015) on each data subset. Significance was determined using Satterwaite’s degrees of freedom method (Satterthwaite, 1946) implemented using the *lmerTest* package (Kuznetsova et al., 2017). The distribution of each alpha diversity metric was assessed graphically and transformed towards normality as appropriate.

The following variables were included as main effects: age; sex; habitat-type: coniferous/mixed; distance to edge, i.e. distance from the nest to the nearest woodland edge; maximum brood size of nest; first-egg lay date of nest. In the *all birds* global models, all pairwise interactions between these main effects (except for lay date × brood size, which were strongly colinear; sex also excluded because it was unavailable for nestlings) were included. In the *adults only* global models, all pairwise interactions were included, except for lay date × brood size and any habitat or age interactions, due to an imbalance in these factors in the dataset, (see table S1). Backwards difference coding was used, instead of the default dummy coding, when fitting the age variable which allowed us to compare each age group to the previous age group sequentially, rather than to a single reference level. This contrast scheme gives sum contrasts, so coefficients reflect main effects rather than marginal effects. Woodland site, nest ID and individual ID were fit to model as nested random intercepts (in the form woodland site/nest ID/individual ID), to control for non-independence. Bird ID was dropped from the *all birds* Chao1 model and woodland site dropped from both adult diversity models due to singular fit warnings.

##### Phylum-level-Relative-Abundance-GLMMs

The two most abundant phyla (Proteobacteria, mean ± SE: 41.6% ± 2.1%; Firmicutes, mean ± SE: 36.6% ± 2.2%) were modelled separately to determine which variables correlated with changes in the phyla relative abundance, in order to develop a broad sense of how the microbial community changed with varying ecological factors. Binomial Generalised Linear Mixed Models (GLMMs) were used. The binomial response variable was ‘phyla abundance’ (sequence reads) weighted by the total number of sequence reads, per sample. These data were also subset into *all birds* and *adults only* subsets, and we used the same model formulas as the alpha diversity models above. The *all birds* global models did not converge so simplified models were refit only including pairwise interactions with age. The age variable was coded using backwards difference coding, as in the alpha diversity models.

#### Model averaging

Model averaging was performed using the MumIn package (Bartoń, 2020) as outlined by Grueber et al. (2011), on the alpha diversity models and the phylum level relative abundance models. In each case, a full model was fit using *lmer* or *glmer*, variables were standardised using the arm package (Gelman et al., 2021) and then a model set generated with the dredge command. The top model set was selected from the complete model set, with models within 2AICc of the top model considered part of this top model set. If multiple models were within the 2AICc cut-off, these were averaged, otherwise the single top model was reported. The input variables were standardised using the Gelman procedure (Gelman, 2008) centring all regression inputs on zero and dividing by 2SDs.

#### Model checking and plotting

Model diagnostics for the alpha diversity and phylum level relative abundance models were checked with *DHARMa* (Hartig, 2019). *DHARMa* simulates data based on the model provided and is a better validator than simple residual vs fitted data plots for mixed effects models. Simulated residual plots were made for each of the models in the top model set. In cases where the simulated residuals showed a clear pattern, models were interpreted cautiously. The *adults only* phyla level models showed overdispersion so these models were refit with an observation level random term following Harrison (2015).

All plots were created using ggplot2 (Wickham, 2016). Alpha diversity and phylum-level results were plotted using ‘JTools’ and ‘Interaction’ (Long, 2019, 2022) packages, which are based on ggplot2. The model estimates for the top model in each model set were plotted, not the model averaged results. These plots used partial residuals, which allow the plotting of data accounting for the variables other than the predictor variable of interest and can be better at showing relationships between variables in multivariate mixed models than plotting the raw data. Plots of the raw data vs the response variable were also made to verify the robustness of model results.

##### Beta-diversity

Beta-diversity measures the dis-similarity between two or more communities, and in our analyses consisted of the pairwise distances between all samples. Beta-diversity was analysed using a compositional approach, which accounts for the proportional nature of high-throughput sequencing data (Gloor et al., 2016; Gloor & Reid, 2016). Low prevalence taxa, i.e. those with less than one copy in 5% of samples, were excluded to reduce possible contaminants and sequencing artefacts (Bokulich et al., 2013). The filtered dataset was centre-log ratio transformed and then the Euclidean (or Aitchison) distance between samples was calculated (Aitchison, 1983). The beta-diversity of samples was tested using PERMANOVA (*adonis2* function from the vegan package (Oksanen et al., 2019)) after checking variables for homogeneity of dispersion. Models were fit using the ‘margin’ option which tests the marginal effect of each variable while accounting for the other variables in the model. Nest was used as a blocking factor to control for the non-independence of samples and sequence plate was included as a fixed effect to account for batch effects. Predictor variables were centred and scaled. For the *all birds* dataset two models were run: one with only non-interacting fixed effects, and another model which included the main interaction of interest (age × habitat), as PERMANOVA cannot calculate the marginal effect of fixed effects from the interactions in the model. For the *adults only* dataset a single model was run with only non-interacting fixed effects. No variable, except habitat, had homogenous dispersion when all birds were considered (table S2), so significant PERMANOVA results could reflect differences in group variance rather than differences in group means, or could reflect differences in both group variance and group means (Anderson & Walsh, 2013). When adult birds were considered separately age, sex and habitat all had homogenous dispersions (table S3). Beta diversity was visualised using PCA plots of the (pairwise) Euclidian distance between samples, and plotted samples according to their first and second principal component values.

#### Repeatability

The rptR package (Stoffel et al., 2019) was used to decompose the components of variance underlying alpha diversity in nestlings measured across D8 and D15, and to examine the extent to which consistent differences among nestlings were confounded by the fixed effects of age and habitat, and the random effects of nest or woodland site. Specifically, repeatability of Shannon and Chao1 diversity was calculated at the individual and group level in the following ways: (i) unadjusted repeatability for woodland site; (ii) unadjusted repeatability for nest; (iii) unadjusted individual repeatability; (iv) individual repeatability adjusted for the fixed effects of age × habitat, (v) repeatability of individual and nest adjusted for age × habitat; and (vi) repeatability of individual, nest and woodland site adjusted for age × habitat. Adjusted repeatability is repeatability controlling for the fixed effect, where any variance explained by fixed effects is excluded from the total variance used in the repeatability estimate (Nakagawa & Schielzeth, 2010). Repeatability was adjusted for age, habitat and their interaction because age × habitat was significant in the nestling only models of alpha diversity (table S4.). Repeatability for phyla-level relative abundance was also calculated in the same manner, adjusting for different fixed effects, according to a nestling only binomial model (see table S5). Singleton nestling measurements were included to improve estimates (Martin et al., 2011). Repeatability analyses were restricted to nestlings because no repeat measures were available for the adults.

## Results

### Alpha diversity

There was no relationship between Shannon diversity and age (table S6a). Chao1 diversity was lower in D15 than in D8 nestlings (−0.35±0.15, p=0.02; table S6b), and there was no difference between D15 nestlings and adult birds (−0.02±0.18, p=0.91; table S6b, figure 1). The effect of habitat on alpha diversity depended on age (table S6a, b) – the decrease in Chao1 diversity from D8 to D15 was more pronounced in mixed/deciduous than in coniferous habitats (habitat × age: −0.73±0.32, p=0.025; table S6b, figure 1). As nest box distance from woodland edge increased, so did both Shannon and Chao1 diversity, regardless of age (Shannon: 0.21±0.1, p=0.035; Chao1: 0.45±0.2, p=0.023; table S6, figure 2). Neither brood size nor lay date predicted alpha diversity (table S6). None of the variables (age, sex, habitat, distance to edge, brood size or lay date) predicted alpha diversity when adults were analysed separately (table S7).

**Figure 1.**
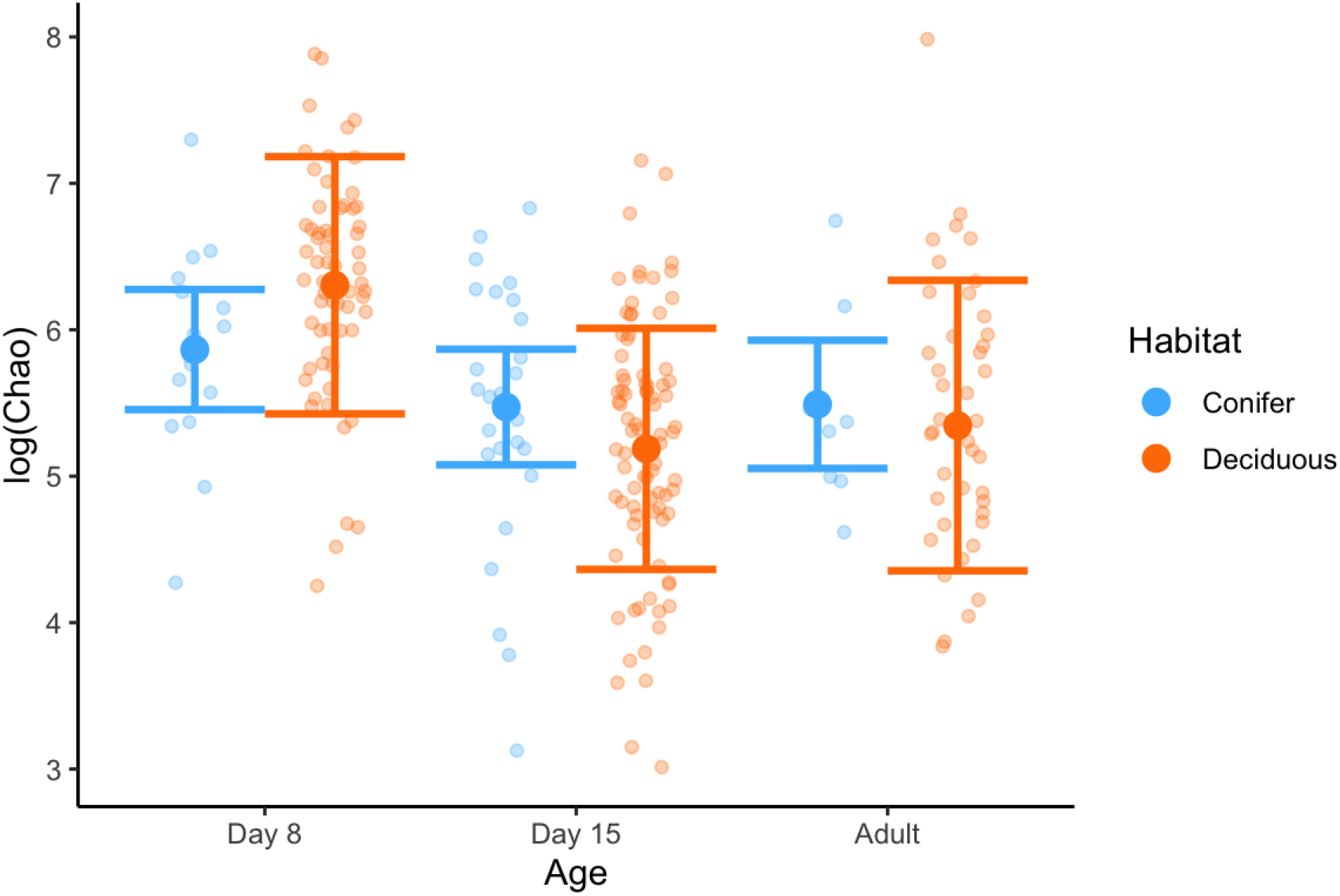
Partial residuals plot for Chao1 diversity regressed on age across habitats (*all birds* model), means with 95% confidence intervals.

**Figure 2.**
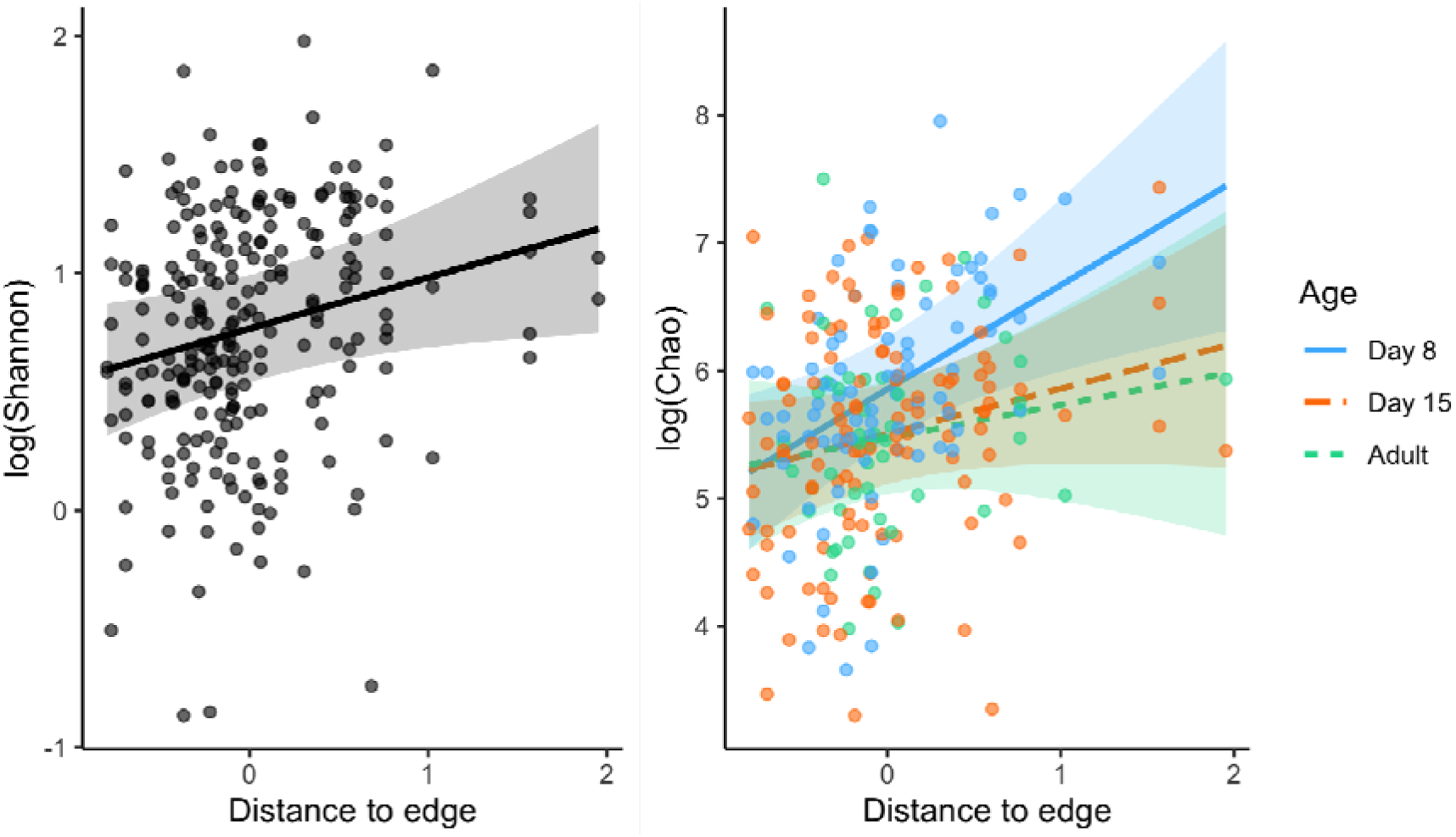
Fitted line on partial residuals plot with 95% confidence intervals for logged diversity regressed on distance to edge from the *all birds* diversity models. Distance to edge variable was centred and scaled so some values are below zero. Interaction trend in chao model so slopes plotted separately by age.

### Beta diversity

Beta diversity differed across age (table 1a) where community composition became more similar among individuals with age (figure 3a). Beta diversity varied with habitat as a main effect (table 1a) and when interacting with age (table 1b). Visual inspection of the PCA plot suggests that the distinction between the ages was most pronounced in the mixed/deciduous woodland (figure S1), though dispersion tests indicate this result could be due to a difference in group dispersions, group means or both (table S2). Beta diversity varied with brood size and distance to edge, and there was also a tendency for beta diversity to vary with lay date (table 1a). When adults were analysed separately, there was no difference in beta diversity between mature and immature birds or between sexes (table 1c), and brood size was the only factor to predict beta diversity (table 1).

**Figure 3.**
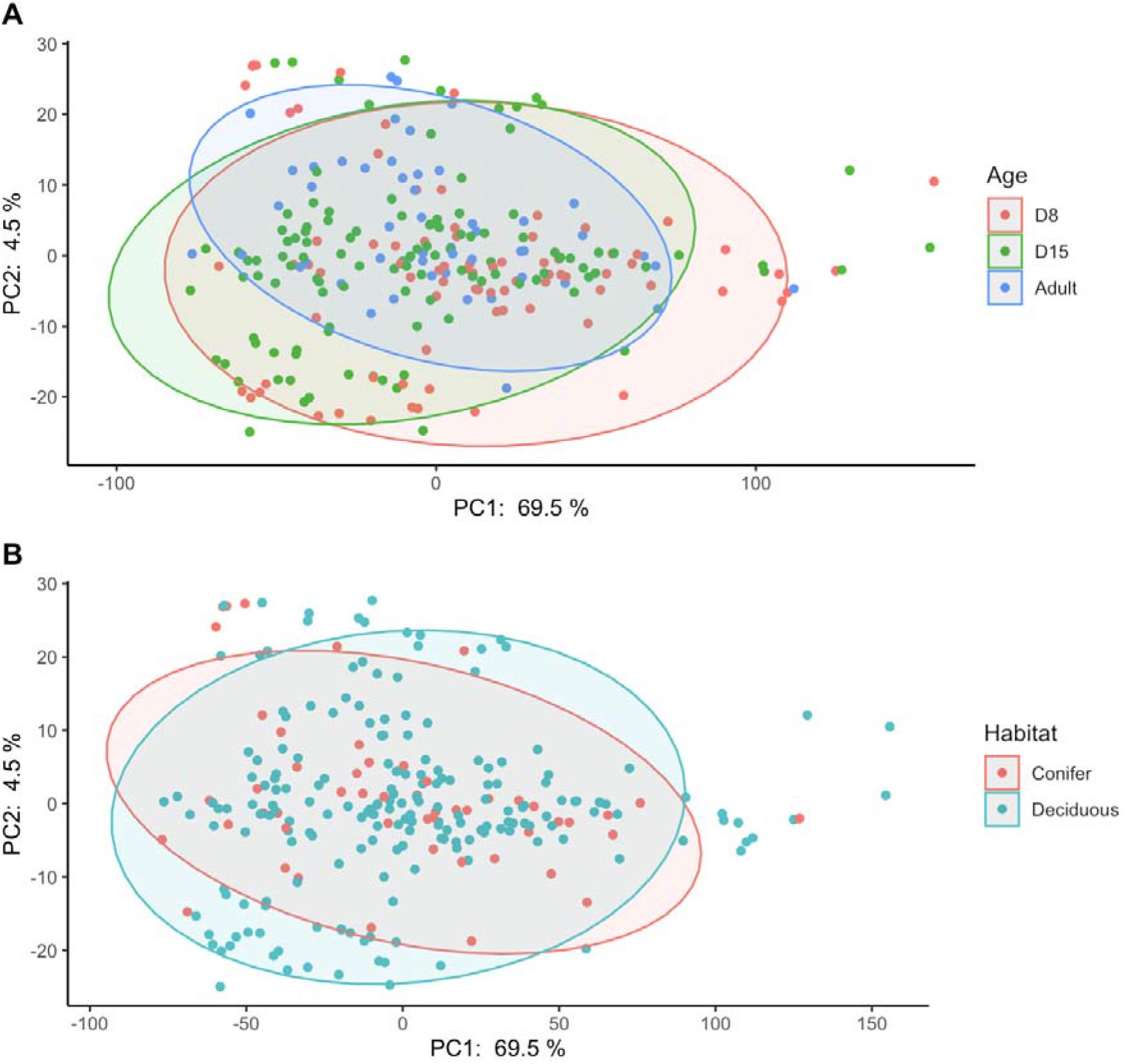
PCA plot of beta diversity with ellipses around (A) age groups, and (B) habitat types.

**Table 1.** Beta-diversity PERMANOVA. Results are marginal effects and significance is calculated from 999 permutations. (a) model with only main effects, (b) model which additionally includes age × habitat interaction term (see methods for explanation) and (c) adults only model. Cells blank where variable not tested.

|  | (a) All birds |  |  |  | (b) All birds (with interaction) |  |  |  | (c) Adults only |  |  |  |
| --- | --- | --- | --- | --- | --- | --- | --- | --- | --- | --- | --- | --- |
| Independent variables | SS | R<sup>2</sup> | F | P | SS | R<sup>2</sup> | F | P | SS | R<sup>2</sup> | F | P |
| Sex |  |  |  |  |  |  |  |  | 212.96 | 0.02 | 0.82 | 0.83 |
| Age | 1062.07 | 0.02 | 2.31 | 0.001 |  |  |  |  | 271.04 | 0.02 | 1.05 | 0.63 |
| Habitat | 307.08 | 0.01 | 1.34 | 0.013 |  |  |  |  | 290.32 | 0.02 | 1.12 | 0.44 |
| Lay date | 423.03 | 0.01 | 1.84 | 0.07 | 419.75 | 0.01 | 1.83 | 0.08 | 323.59 | 0.02 | 1.25 | 0.13 |
| Brood size | 318.83 | 0.01 | 1.39 | 0.006 | 305.34 | 0.01 | 1.33 | 0.018 | 223.30 | 0.02 | 0.86 | 0.004 |
| Distance to edge | 496.18 | 0.01 | 2.16 | 0.001 | 502.35 | 0.01 | 2.19 | 0.001 | 327.04 | 0.02 | 1.26 | 0.34 |
| Sequence plate | 3653.08 | 0.06 | 3.97 | 0.001 | 3634.14 | 0.06 | 3.96 | 0.001 | 1462.09 | 0.11 | 1.88 | 0.009 |
| Age × Habitat | | | | | 594.52 | 0.01 | 1.30 | 0.015 | | | | |

### Phylum level relative abundance

D8 nestlings had higher Proteobacteria abundance than D15 nestlings, who in turn had lower Proteobacteria abundance than adults (table 2a and figure 4A). The main effect of age also interacted with habitat. In both habitats, Proteobacteria decreased with nestling age (i.e. between D8 and D15), but this effect was stronger for birds developing in mixed/deciduous compared to coniferous habitats, mirroring the pattern above for alpha diversity (table 2, figure 4A, 4B). Adult birds had greater abundances of Proteobacteria than D15 birds, and this effect was especially pronounced in mixed/deciduous woodlands compared to conifer woodlands (table 2b; figure 4A). We note that although the age-habitat interactions are supported by the model, they are not strongly supported by plots (Fig 5A and B) and should be interpreted with caution. Brood size was negatively related to the abundance of Proteobacteria in nestlings, especially for D15, but not for adults; the opposite was the case for Firmicute abundance (table 2a and 2b; figure 4C and 4D). There was no effect of distance to edge on phyla level abundance. Proteobacteria abundance was negatively related to lay date in D8 nestlings, and positively in D15 nestlings and in adults. All of the patterns described for Proteobacteria were found for Firmicutes in the inverse directions (figure 4, table 2). Only lay date predicted phyla level relative abundance in the *adults only* models. Proteobacteria increased with lay date (1.5±0.5, p=0.006), but did not relate to Firmicutes (−0.7±0.6, p=0.223; table S8, figure 5).

**Table 2.** Phylum level, binomial model results for the *all birds* (nestlings and adults) subset. Only a single model was retained in the top model sets (<2AICc) so results are not model averaged. The reference level for habitat is ‘coniferous’ which is compared here to ‘mixed/deciduous’. The age variable is backwards difference coded (see methods for explanation).

| Independent variables | (a) Proteobacteria |  |  |  | (b) Firmicutes |  |  |  |
| --- | --- | --- | --- | --- | --- | --- | --- | --- |
|  | Estimate | S.E. | z | P | Estimate | S.E. | z | P |
| (Intercept) | 0.16 | 0.67 | 0.24 | 0.81 | -2.17 | 0.70 | -3.12 | 0.002 |
| Age (D15/D8) | -1.52 | 0.00 | -342.63 | <0.001 | 1.30 | 0.00 | 315.88 | <0.001 |
| Age (Adult/D15) | 2.45 | 0.31 | 7.88 | <0.001 | -2.94 | 0.35 | -8.47 | <0.001 |
| Habitat | -0.08 | 0.72 | -0.12 | 0.91 | 0.08 | 0.89 | 0.09 | 0.93 |
| Brood size | -0.72 | 0.20 | -3.58 | <0.001 | 0.97 | 0.22 | 4.36 | <0.001 |
| Distance to edge | 0.66 | 0.41 | 1.62 | 0.11 | 0.12 | 0.48 | 0.25 | 0.81 |
| Lay date | 0.49 | 0.44 | 1.11 | 0.27 | -0.38 | 0.53 | -0.70 | 0.48 |
| Age (D15/D8) × Habitat | -0.87 | 0.01 | -129.32 | <0.001 | 1.92 | 0.01 | 302.68 | <0.001 |
| Age (Adult/D15) × Habitat | 1.90 | 0.84 | 2.26 | 0.024 | -1.67 | 0.91 | -1.84 | 0.07 |
| Age (D15/D8) × Brood size | -1.19 | 0.01 | -174.16 | <0.001 | 1.36 | 0.01 | 211.54 | <0.001 |
| Age (Adult/D15) × Brood size | 1.90 | 0.61 | 3.14 | 0.002 | -1.28 | 0.66 | -1.93 | 0.054 |
| Age (Day15/Day8) × Distance to edge | -0.63 | 0.01 | -115.05 | <0.001 | 0.73 | 0.01 | 129.92 | <0.001 |
| Age (Adult/D15) × Distance to edge | -0.66 | 0.65 | -1.01 | 0.31 | 0.34 | 0.71 | 0.47 | 0.64 |
| Age (D15/D8) × Lay date | 1.77 | 0.01 | 319.66 | <0.001 | -1.67 | 0.01 | -309.36 | <0.001 |
| Age (Adult/D15) × Lay date | 0.67 | 0.79 | 0.85 | 0.40 | -0.34 | 0.84 | -0.40 | 0.69 |

**Figure 4.**
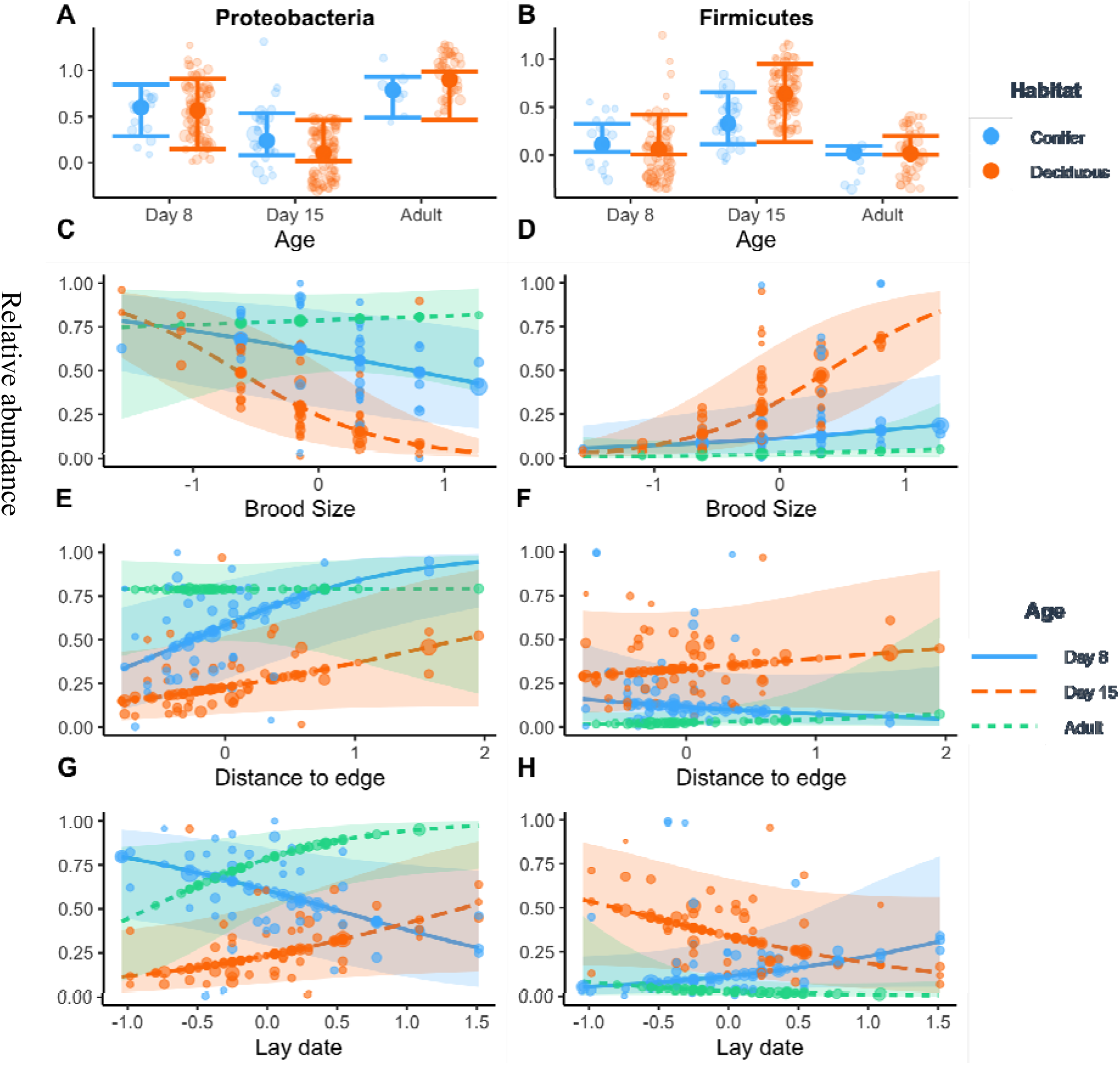
Partial residual plots for relative abundance of Proteobacteria (left column) and Firmicutes (right column) for *all birds* phyla-level models showing means with 95% confidence intervals. Point size reflects the weight of that observation, i.e. the ‘number of sequence reads’ of the sample. Some points may appear out of range due to ‘jitter’ function which separates overlapping points.

**Figure 5.**
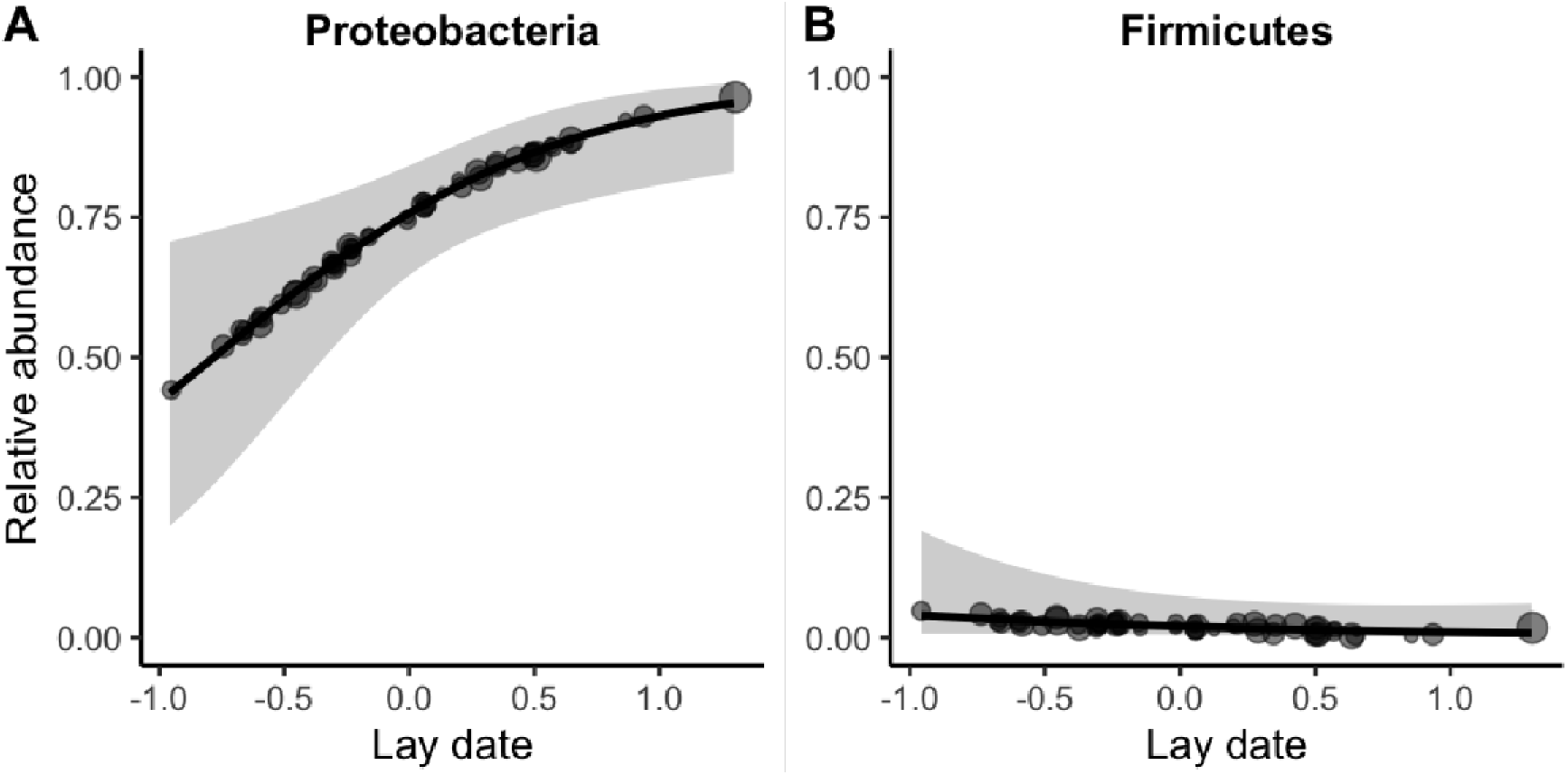
Partial residual plots for relative abundance of Proteobacteria and Firmicutes against lay date (adult birds models), means with 95% confidence intervals.

### Nestling, nest and woodland site repeatability

Microbiota were not repeatable across sites (model 1’s, table S9) but were repeatable across nests for Shannon, Chao1, Proteobacteria and Firmicutes abundance (R values from model 2’s, respectively: 0.399; 0.421; 0.191; 0.261. See table S9 for credibility intervals and further details). Individual gut microbiota were not significantly repeatable for Shannon diversity, but was for Chao1 diversity, Proteobacteria abundance and Firmicutes abundance, when unadjusted for any other effects (R values from model 3s, respectively: 0.378, n.s.; 0.46; 0.174; and 0.248). After controlling for fixed effects, individual repeatabilities for Shannon and Chao1 diversity, but not for Proteobacteria or Firmicutes relative abundance, were significant (R values from model 4’s in table S9 respectively: 0.403; 0.50; 0.144, n.s.; 0.165, n.s.). However, individual repeatabilities for all four gut microbiota metrics approached zero when controlling for nest site as a random effect, which itself was repeatable in all four variables (see R values from model 5’s for both individual and nest in table S9), indicating that the majority of microbiota variability in nestlings was driven by nest identity, not individual differences. The addition of woodland site as a random effect suggested that up to a quarter of the nest site repeatability may have been driven by differences among sites (table S9), though the credibility intervals for woodland site repeatability overlapped zero in all four model 6’s.

Given these repeatability results, we decided to carry out post-hoc analyses of how parent traits correlated with offspring traits. We found that nestling microbiome was predicted only by the mother’s Shannon diversity (table S10a-d). Specifically, female Shannon diversity negatively predicted offspring Shannon diversity (figure S2). None of mother’s Chao1, Proteobacteria abundance or Firmicutes abundance, or any of the father’s microbiome traits predicted the corresponding nestling trait (table S10e-h).

## Discussion

We found that environmental links with the gut microbiota were dependent on age, pointing to differential sensitivity of gut microbiota across life stages in response to habitat type, local environmental gradients and life history traits (table 3), which is consistent with results from human studies (Claesson et al., 2011). Notably, life history, but not environmental effects predicted gut microbiota variation in adults; whereas nestling gut microbiota appeared sensitive to a range of both environmental and life history variations (table 3). Repeated measures showed that although nestlings appeared to show consistent individual differences in gut microbiota diversity and phylum-level abundance, this was explained entirely by the nest in which they developed. The post-hoc parent-offspring analyses found the microbiome traits of nestlings were not typically correlated with their parents’ traits, although nestlings’ Shannon diversity was negatively predicted by their mother’s Shannon diversity. We discuss differential gut microbiota sensitivities to environmental and life history variation across age groups and place these correlative findings in the context of future directions for understanding the ecological and evolutionary consequences of these patterns in wild systems.

**Table 3.**
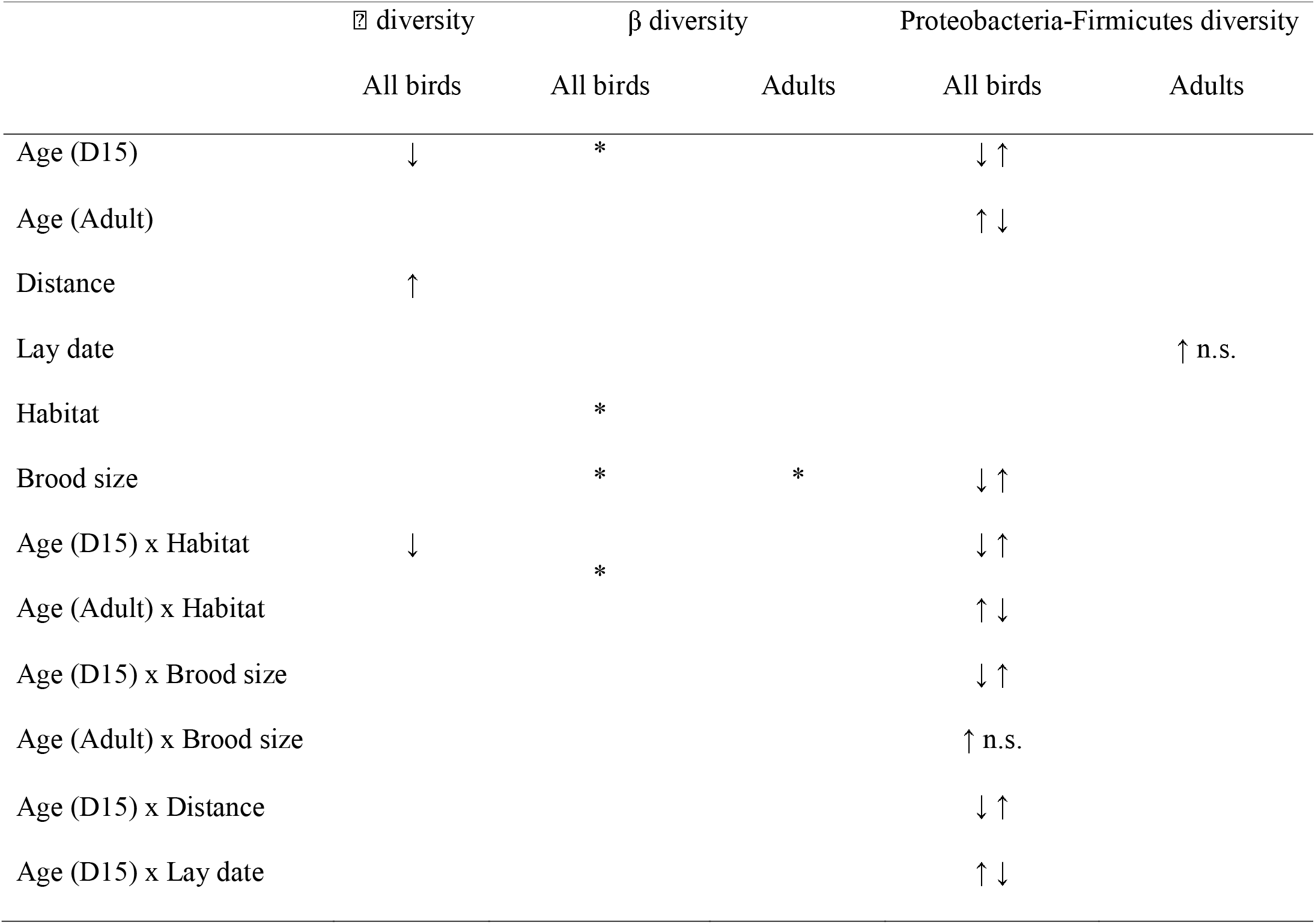
Summary of results. Arrows show the direction of the effect on the microbiome trait (not applicable for β diversity as effects do not have directionality). Paired arrows indicate the direction of effect for Proteobacteria and Firmicutes, respectively. Asterisks show significant results for beta diversity. ‘Coniferous’ is the reference level for habitat and bracketed term with ‘Age’ indicates age level which is being compared to the proceeding level.

### Environment

Only nestling, not adult, gut microbiota varied in response to woodland type (table 3). This is not surprising given microbial community assembly theory (Costello et al., 2012; Coyte et al., 2021) where source microbes vary according to the external environment (e.g. diet, nesting material, habitat) (C.-Y. Chen et al., 2020; Goodenough et al., 2016; Goossens et al., 2022; Koenig et al., 2010) and compete for available niches as the host physiological environment develops (Costello et al., 2012). Diet has a strong seasonal component during this time, as the availability of food items changes throughout the breeding season and varies with habitat type, habitat quality and parental preferences (Wilkin et al., 2009). Regardless of age, the alpha diversity of the microbiota was higher for birds in nests further from the edge of the woodland (figure 2), with a trend for the strength of this association to diminish with age, which we hypothesise could be due to differences in the exposure to microbes via nesting materials (Goodenough et al., 2016) and diet (G. Davidson et al., 2020) during a time when the microbiota are particularly sensitive to environmental variation. Similarly, D8 nestlings were especially sensitive to the woodland habitat (mixed/deciduous or coniferous), whereas the effect of woodland on alpha diversity was not present in adults, nor did it differ between adults and D15 nestlings, suggesting that the effects of habitat variation on alpha diversity stabilised early in development. However, for phylum-level abundance, this stability was not observed until adulthood. Future studies should collect microbiota community data and with contemporary DNA metabarcoding data across environmental gradients to determine whether the microbiota covaries with diet across habitats.

### Life history and age

Lay date was positively correlated to Proteobacteria in D15 nestlings and in adults, but negatively in D8 nestlings, while the exact reverse was true for Firmicute abundance. The abundance of caterpillars, the primary food source for developing great tits, peaks mid breeding season, the precise timing of which could affect the gut microbiota directly, or indirectly through the effects of food availability on stress, immunity or growth (Hooper et al., 2012; Potti et al., 2002; Stothart et al., 2019). Indeed, nestlings with an early or late lay-date may experience stress from lower food availability and higher predation pressure (Naef-Daenzer et al., 2001), and experimentally manipulated glucocorticoids have been reported to affect nestling bird gut microbial diversity (Noguera et al., 2018). Furthermore, the effect of lay date on stress may depend on nestling age because of the increasing nutritional demands of older nestlings, so that young nestlings from early-nesting parents could face similar conditions to old nestlings from late-nesting parents. If, hypothetically, the positive correlation observed between adult Proteobacteria and lay date was driven by stress, this could explain why the effect of lay date on phylum-level abundance was in opposing directions for D8 and D15 nestlings: D15 nestlings from early nests may have been equally stressed to D8 nestlings from late nests (figure 4G).

Brood size also correlated with microbiota variation, which we suggest is likely due to larger brood sizes leading to higher stress as a consequence of increased sibling competition (Neuenschwander et al., 2003) and lower food availability (Smith et al., 1988). The effect of brood size on microbiota was especially pronounced for D15 birds, when demand for food is highest and sibling-sibling conflict greatest. It is possible that larger brood sizes simply have a different reservoir of bacteria due to higher levels of provisioning. Cross fostering experiments that manipulate the size of broods, and compares broods with nestlings from multiple source nests against broods with nestlings from a single source, would test the relative importance of this reservoir and competition hypotheses.

Contrary to our predictions, in the *adults only* dataset, we did not detect any effects of lay date on gut microbiota, nor any sex-dependent interactions, despite lay date being strongly determined by female condition (Brown & Brown, 1999). Lay date could also potentially affect the adult gut microbiota indirectly, via increased glucocorticoids related to provisioning effort (Bonier et al., 2011), because provisioning the young is more challenging outside of the peak of food abundance. Although stress hormones have been reported to alter gut microbiota in reproductive female lizards (*Sceloporus undulatus*), these effects were only observed at specific reproductive stages (MacLeod et al., 2022). In the current study, adults were sampled for gut microbiota near the end of their reproductive cycle (i.e. when chicks were near fledging), and therefore it is possible we missed key reproductive time points where gut microbiota were sensitive to physiological influences associated with lay date. Nevertheless, brood size, which may also reflect parental quality (Pettifor et al., 2001) and/or higher glucocorticoid demands associated with provisioning (Bonier et al., 2011), predicted gut microbiota community composition in adults of both sexes. Overall, these findings highlight a potentially exciting area of research on reproductive-dependent physiology and parental care, and its interaction with the function of the gut microbiome. Brood size manipulation in conjunction with glucocorticoid measurements would determine whether the effect of brood size on the microbiota was related to parental quality or glucocorticoid levels.

### Repeatability

The repeatability analysis revealed the importance of the immediate nest environment in structuring the microbiota, whereas the larger-scale effect of woodland site was negligible. Human parents provide their offspring with beneficial microbes acquired over their own lifespan and hence provide their offspring a fitness advantage (Sprockett et al., 2018). These microbes prime them for their environment during early life and are particularly influential due to priority effects, where the timing of colonisation influences future species interactions (Sprockett et al., 2018). Notably, individual-level repeatability of microbial diversity was zero when nest identity was controlled, where individual’s gut microbiota were not following individual trajectories but were converging on their nestmates. The importance of the nest in determining microbiota diversity and structure has been highlighted in this species and others previously (Campos-Cerda & Bohannan, 2020; Teyssier, Lens, et al., 2018). If genetics were an important driver of the microbiota in nestlings we would expect to see some level of repeatability, even when controlling for nest as siblings are expected to share only 50% of their DNA on average. These results point to considerable plasticity in microbiota assembly during early development, and given the negligible evidence for positive correlation between any parent and offspring microbiome trait, is further support to the greater influence of local environmental factors, most likely linked to local food availability, over intrinsic or host factors, in avian microbiota (Song et al., 2020). Despite these findings, the negative correlation for the mother’s Shannon diversity raises the possibility of a maternal effect on the gut microbiota, which has been demonstrated in wild North American red squirrels (Tamiasciurus hudsonicus) using a quantitative genetic approach (Ren et al., 2017).

## Conclusions

Studies of environmental associations with the gut microbiome in natural systems typically focus on a single ecological feature (e.g. habitat type, season), when in fact multiple interacting ecological factors likely shape the gut microbiome simultaneously. Our results point to links between different aspects of the gut microbiota and multiple aspects of the environment and life history variation: the local nest, sibling interactions, location within woodland plots, the woodland plots themselves, and potentially changes in food availability during the season. These effects are to be expected since many of the factors involved are fundamental drivers of ecological processes within populations, and yet notably, it was the natal, as opposed to the adult gut microbiota that was most sensitive to ecological variation, pointing to a high degree of developmental plasticity. Unpicking the mechanisms, causes and consequences of gut microbiome variation among wild populations is an exciting avenue of future research that remains largely unexplored.

## Acknowledgements

We thank Iván de la Hera, for assistance with fieldwork, and Niamh Wiley for her help with DNA extractions and sequencing. We also thank Paul Cotter, Fiona Crispie and Laura Finnegan from the Teagasc Sequencing facility for their role in relation to the 16S rRNA sequencing. We thank Fiona Fouhy, Orla O’Sullivan and Calum Walsh for their advice on bioinformatics. We are grateful to James Nichols, Ally Phillimore and Sarah Knowles for advice on molecular methodology, and to Caroline McKeon, Darío Fernández-Bellon and Enrico Pirotta for helpful discussions on statistical analyses.

## Data accessibility statement

Metadata is available at DataDryad https://datadryad.org/stash/dataset/doi:10.5061/dryad.bk3j9kd9g (Davidson, et al. 2020). Sequence data are available in the European Nucleotide Archive under access number PRJEB42330, and ERS5506097 – ERS5506338.

