## Supplemental_1 for "Individual variation in the avian gut microbiota: the influence of host state and environmental heterogeneity"

**Electronic Supplementary Material**

| Table S1. Breakdown of sampling. Fewer coniferous birds were sampled due to lower population density of conifer compared to mixed deciduous forests. | | | |
| --- | --- | --- | --- |
|  | **Day-8** | **Day-15** | **Adult** |
| Conifer | 16 | 27 | 8 (Female = 3; Male = 5) |
| Deciduous | 65 | 87 | 43 (Female = 21; Male = 22) |

Table S2. Betadisper results for each variable in the *all birds* PERMANOVA models. Significant results indicate violation of homogeneity of variance assumption of PERMANOVA.

|  | **Sum Sq.** | **Mean Sq.** | **P** |
| --- | --- | --- | --- |
| Age | 107 | 53.5 | 0.025 |
| Habitat | 5.7 | 5.7 | 0.55 |
| Lay date | 1047.2 | 38.8 | <0.001 |
| Brood size | 220 | 36.7 | 0.025 |
| Distance to edge | 2515.5 | 41.9 | <0.001 |
| Sequence plate | 541.7 | 135.4 | <0.001 |

Table S3. Betadisper results for each variable in the adult PERMANOVA models. Significant results indicate violation of homogeneity of variance assumption of PERMANOVA.

|  | **Sum Sq.** | **Mean Sq.** | **P** |
| --- | --- | --- | --- |
| Age | 3.18 | 3.18 | 0.562 |
| Sex | 13.81 | 13.8 | 0.278 |
| Habitat | 6.93 | 6.93 | 0.4196 |
| Lay date | 1328 | 60.4 | <0.001 |
| Brood size | 407.7 | 81.5 | <0.001 |
| Distance to edge | 1559.5 | 42.15 | <0.001 |
| Sequence plate | 142.8 | 47.6 | 0.002 |

Table S4. Alpha diversity, model averaged results for the *nestling only* subset, for the a) Shannon index and b) Chao1 index. **†**Adjusted standard errors.

|  | (a) **Shannon** | | | | (b) **Chao1** | | | |
| --- | --- | --- | --- | --- | --- | --- | --- | --- |
|  | **Estimate** | **SE†** | **z value** | **P** | **Estimate** | **SE†** | **z value** | **Pr(>\|z\|)** |
| (Intercept) | 0.794 | 0.145 | 5.475 | <0.001 | 0.794 | 0.145 | 5.475 | <0.001 |
| Age | -0.146 | 0.090 | 1.627 | 0.10 | -0.146 | 0.090 | 1.627 | 0.10 |
| DistanceToEdge | 0.236 | 0.130 | 1.819 | 0.07 | 0.236 | 0.130 | 1.819 | 0.07 |
| Habitat | 0.112 | 0.215 | 0.522 | 0.60 | 0.112 | 0.215 | 0.522 | 0.60 |
| Age x habitat | -0.387 | 0.191 | 2.028 | 0.04 | -0.387 | 0.191 | 2.028 | 0.04 |
| Habitat x Distance | 0.393 | 0.263 | 1.494 | 0.14 | 0.393 | 0.263 | 1.494 | 0.14 |
| Lay date | 0.144 | 0.145 | 0.989 | 0.32 | 0.144 | 0.145 | 0.989 | 0.32 |
| Brood size | -0.104 | 0.120 | 0.869 | 0.38 | -0.104 | 0.120 | 0.869 | 0.38 |
| Age x DistanceToEdge | -0.074 | 0.165 | 0.449 | 0.65 | -0.074 | 0.165 | 0.449 | 0.65 |

Table S5. Binomial model results for phylum level relative abundance for *nestling only* subset. These results informed the fixed effects used in the repeatability analyses. **†** Adjusted standard errors.

| **Independent variables** | **Estimate** | **SE†** | **z** | **P** | **Estimate** | **SE†** | **z** | **P** |
| --- | --- | --- | --- | --- | --- | --- | --- | --- |
|  | **(a) Proteobacteria** | | | | **(b) Firmicutes** | | | |
| (Intercept) | -0.80 | 0.76 | 1.05 | 0.292 | -0.63 | 0.85 | -0.74 | 0.46 |
| Age | -1.24 | 0.01 | 264.95 | <0.001 | 0.97 | 0.00 | 225.74 | <0.001 |
| Habitat | 1.14 | 1.12 | 1.02 | 0.307 | -2.74 | 1.43 | -1.91 | 0.056 |
| Brood size | 0.27 | 0.01 | 22.18 | <0.001 | -1.45 | 0.01 | -100.06 | <0.001 |
| Distance to edge | 0.41 | 0.56 | 0.73 | 0.463 | 1.20 | 0.78 | 1.54 | 0.125 |
| Lay date | 0.40 | 0.66 | 0.62 | 0.538 | -0.59 | 0.91 | -0.65 | 0.515 |
| Age x habitat | -2.63 | 0.01 | 242.53 | <0.001 | 5.01 | 0.01 | 476.80 | <0.001 |
| Age x brood size | -0.92 | 0.01 | 116.50 | <0.001 | 1.91 | 0.01 | 249.46 | <0.001 |
| Age x distance to edge | 1.35 | 0.01 | 110.43 | <0.001 | -3.75 | 0.01 | -281.08 | <0.001 |
| Age x lay date | 2.06 | 0.01 | 331.83 | <0.001 | -1.86 | 0.01 | -308.61 | <0.001 |
| Habitat x brood size | -8.32 | 0.05 | 162.79 | <0.001 | 14.98 | 0.06 | 234.90 | <0.001 |
| Habitat x distance | 3.15 | 1.25 | 2.52 | 0.012 | -4.83 | 1.62 | -2.99 | 0.003 |
| Brood size x distance to edge | 2.22 | 0.04 | 63.62 | <0.001 | -5.16 | 0.05 | -112.53 | <0.001 |
| Distance to edge x lay date | 1.32 | 1.63 | 0.81 | 0.42 |  |  |  |  |
| Habitat x lay date | -2.15 | 2.37 | 0.91 | 0.365 |  |  |  |  |

| Table S6. Alpha diversity, model averaged results for the *all birds* subset, for the a) Shannon index and b) Chao1 index. Cells are blank where the variable was not retained following the model selection procedure. **†** Adjusted standard errors. | | | | | | | | |
| --- | --- | --- | --- | --- | --- | --- | --- | --- |
|  | **(a)   Shannon** | | | | **(b)   Chao1** | | | |
| **Independent variables** | **Estimate** | **SE†** | **z value** | **P** | **Estimate** | **SE†** | **z value** | **P** |
| (Intercept) | 0.77 | 0.12 | 6.65 | <0.001 | 5.61 | 0.19 | 29.85 | <0.001 |
| Age (D15/D8) |  |  |  |  | -0.35 | 0.15 | 2.33 | 0.02 |
| Age (Adult/D15) |  |  |  |  | -0.02 | 0.18 | 0.12 | 0.91 |
| Distance to edge | 0.21 | 0.10 | 2.11 | 0.035 | 0.45 | 0.20 | 2.27 | 0.023 |
| Lay date | 0.08 | 0.11 | 0.73 | 0.46 |  |  |  |  |
| Age(D15-D8) × Distance-to- edge |  |  |  |  | -0.46 | 0.28 | 1.68 | 0.09 |
| Age(Adult-D15) × Distance-to-edge |  |  |  |  | -0.09 | 0.33 | 0.28 | 0.78 |
| Brood size |  |  |  |  | 0.15 | 0.18 | 0.83 | 0.41 |
| Habitat |  |  |  |  | 0.00 | 0.32 | 0.01 | 0.99 |
| Age(D15/D8) × Habitat |  |  |  |  | -0.73 | 0.32 | 2.24 | 0.025 |
| Age (Adult/D15) × Habitat |  |  |  |  | 0.14 | 0.42 | 0.34 | 0.74 |

| **Table S7.** Alpha diversity linear model results for the *adults only* subset, for a) Shannon index and b) Chao1 index. Results are model averaged mixed model values for retained variables. **†**Adjusted SE | | | | | | | | |
| --- | --- | --- | --- | --- | --- | --- | --- | --- |
|  | **(a)   Shannon** | | | | **(b)   Chao1** | | | |
| **Independent variables** | **Estimate** | **SE†** | **z value** | **P** | **Estimate** | **SE†** | **z value** | **P** |
| (Intercept) | 1.31 | 0.05 | 26.24 | <0.001 | 5.45 | 0.19 | 28.22 | <0.001 |
| Brood size | 0.07 | 0.07 | 0.95 | 0.34 | 0.34 | 0.24 | 1.41 | 0.16 |
| Habitat | -0.07 | 0.09 | 0.76 | 0.45 |  |  |  |  |
| Sex | -0.04 | 0.05 | 0.70 | 0.48 |  |  |  |  |
| Distance to edge |  |  |  |  | 0.24 | 0.23 | 1.02 | 0.31 |
| Age(Mature- Juvenile) |  |  |  |  | -0.36 | 0.34 | 1.09 | 0.28 |
| Lay date |  |  |  |  | -0.16 | 0.24 | 0.66 | 0.51 |

| Table S8. Phylum level results for the *adults only* subset for a) Proteobacteria and c) Firmicutes. Results are model averaged binomial mixed model values for retained variables. **†**Adjusted SE | | | | | | | | |
| --- | --- | --- | --- | --- | --- | --- | --- | --- |
|  | **(a)    Proteobacteria** | | | | **(b)    Firmicutes** | | | |
| **Independent variables** | **Estimate** | **SE†** | **z value** | **P** | **Estimate** | **SE†** | **z value** | **P** |
| (Intercept) | 1.137 | 0.273 | 4.169 | <0.001 | -3.865 | 0.679 | 5.692 | <0.001 |
| Age (Mature- Juvenile) | -0.779 | 0.736 | 1.059 | 0.289 |  |  |  |  |
| Distance to edge |  |  |  |  | 0.689 | 0.603 | 1.142 | 0.253 |
| Lay date | 1.458 | 0.525 | 2.775 | 0.006 | -0.743 | 0.609 | 1.219 | 0.223 |
| Brood size |  |  |  |  | 0.531 | 0.641 | 0.828 | 0.408 |

| Table S9. (a) Adjusted and unadjusted values of individual and group level repeatability of (a) nestling alpha diversity (logged), with confidence intervals in brackets. Fixed effects where included are: Age + Habitat + Age × Habitat; (b) (logit link scale approximation) of nestling phylum level relative abundance. Six models were run for each phyla with stepwise addition of fixed and random effects. Fixed effects where included are: Age + Habitat + Lay date + Brood size + Distance to edge + Age × Habitat + Age × Lay date + Age × Brood size + Age × Distance to edge + Brood size × Habitat + Brood size × Distance to edge. | | | | |
| --- | --- | --- | --- | --- |
| (a) | | | | |
|  | Repeatability (CI) | | |  |
|  | Individual repeatability | Nest | Woodland site | Fixed |
| Shannon |  |  |  |  |
| 1 |  |  | 0.146 (0, 0.345) | No |
| 2 |  | 0.399 (0.185, 0.543) |  | No |
| 3 | 0. 378 (0, 0.423) |  |  | No |
| 4 | 0.403 (0.298, 0.455) |  |  | Yes |
| 5 | 0 (0, 0.227) | 0.423 (0.229, 0.562) |  | Yes |
| 6 | 0 (0, 0) | 0.318 (0.139, 0.493) | 0.11 (0, 0.33) | Yes |
| Chao |  |  |  |  |
| 1 |  |  | 0.127 (0, 0.298) | No |
| 2 |  | 0.421 (0.186, 0.632) |  | No |
| 3 | 0.46 (0.399, 0.48) |  |  | No |
| 4 | 0.498 (0.476, 0.515) |  |  | Yes |
| 5 | 0 (0, 0.207) | 0.499 (0.314, 0.641) |  | Yes |
| 6 | 0 (0, 0) | 0.392 (0.194, 0.576) | 0.114 (0, 0.365) | Yes |
| Proteobacteria |  |  |  |  |
| 1 |  |  | 0.067 (0, 0.145) | No |
| 2 |  | 0.191 (0.091, 0.284) |  | No |
| 3 | 0.174 (0.27, 0.286) |  |  | No |
| 4 | 0.144 (0, 0.254) |  |  | Yes |
| 5 | 0.026 (0, 0.14) | 0.178 (0.046, 0.238) |  | Yes |
| 6 | 0.024 (0, 0.145) | 0.139 (0.022, 0.202) | 0.04 (0, 0.112) | Yes |
| Firmicutes |  |  |  |  |
| 1 |  |  | 0.104 (0, 0.198) | No |
| 2 |  | 0.261 (0.138, 0.358) |  | No |
| 3 | 0.248 (0.085, 0.391) |  |  | No |
| 4 | 0.165 (0, 0.315) |  |  | Yes |
| 5 | 0.037 (0, 0.126) | 0.267 (0.099, 0.338) |  | Yes |
| 6 | 0 (0, 0.124) | 0.188 (0.057, 0.271) | 0.078 (0, 0.181) | Yes |

Table S10. Parent offspring analysis. Offspring microbiome traits regressed on parents (a-d: mother; e-h: father) corresponding microbiome trait, controlling for fixed effects and the random effect of nest. Observations from 57 nestlings from 21 nests (a-d) and 59 nestlings from 22 nests (e-h).

| **Independent variables** | **Estimate** | **Std. Error** | **t value** | **P** |
| --- | --- | --- | --- | --- |
| **(a) Shannon diversity** | | | | |
| Shannon (female) | -1.005 | 0.408 | -2.463 | 0.025 |
| Age | -0.879 | 0.328 | -2.681 | 0.009 |
| Habitat | 1.215 | 0.603 | 2.017 | 0.08 |
| Brood size | -0.477 | 0.399 | -1.196 | 0.24 |
| Lay date | -0.342 | 0.700 | -0.488 | 0.631 |
| **(b) Chao diversity** | | | | |
| Chao1 (female) | -0.226 | 0.464 | -0.486 | 0.632 |
| Age | -0.719 | 0.269 | -2.673 | 0.009 |
| Habitat | 0.654 | 0.654 | 0.999 | 0.341 |
| Brood size | 0.170 | 0.362 | 0.469 | 0.641 |
| Lay date | 0.001 | 0.719 | 0.002 | 0.999 |
| **(c) Proteobacteria RA** | | | | |
| Proteobacteria (female) | 0.316 | 0.897 | 0.353 | 0.724 |
| Age | -0.515 | 0.005 | -98.828 | <0.001 |
| Habitat | 0.340 | 1.367 | 0.249 | 0.804 |
| Brood size | -0.907 | 0.006 | -158.429 | <0.001 |
| Lay date | -2.104 | 1.454 | -1.447 | 0.148 |
| **(d) Firmicutes RA** | | | | |
| Firmicutes (female) | 1.496 | 0.855 | 1.749 | 0.08 |
| Age | -0.515 | 0.005 | -98.831 | <0.001 |
| Habitat | -0.218 | 1.452 | -0.150 | 0.881 |
| Brood size | -0.907 | 0.006 | -158.433 | <0.001 |
| Lay date | -2.325 | 1.346 | -1.728 | 0.084 |
| **(e) Shannon diversity** |  |  |  |  |
| Shannon (male) | 0.765 | 0.564 | 1.355 | 0.189 |
| Age | -0.295 | 0.349 | -0.844 | 0.401 |
| Habitat | 0.547 | 0.965 | 0.567 | 0.596 |
| Brood size | -0.172 | 0.440 | -0.391 | 0.697 |
| Lay date | -0.191 | 0.562 | -0.340 | 0.742 |
| **(f) Chao diversity** |  |  |  |  |
| Chao1 (male) | 0.400 | 0.336 | 1.193 | 0.242 |
| Age | -0.233 | 0.298 | -0.781 | 0.437 |
| Habitat | 0.040 | 0.730 | 0.054 | 0.958 |
| Brood size | 0.247 | 0.374 | 0.660 | 0.512 |
| Lay date | 0.008 | 0.494 | 0.015 | 0.988 |
| **(g) Proteobacteria RA** |  |  |  |  |
| Proteobacteria (male) | -0.840 | 0.894 | -0.940 | 0.347 |
| Age | 1.253 | 0.006 | 227.282 | <0.001 |
| Habitat | 0.314 | 1.146 | 0.274 | 0.784 |
| Brood size | 0.629 | 0.006 | 111.184 | <0.001 |
| Lay date | 1.019 | 1.159 | 0.879 | 0.379 |
| **(h) Firmicutes RA** |  |  |  |  |
| Firmicutes (male) | 0.998 | 0.810 | 1.231 | 0.218 |
| Age | 1.185 | 0.004 | 300.861 | <0.001 |
| Habitat | -0.080 | 0.963 | -0.083 | 0.934 |
| Brood size | 0.628 | 0.004 | 158.484 | <0.001 |
| Lay date | 0.774 | 1.009 | 0.767 | 0.443 |

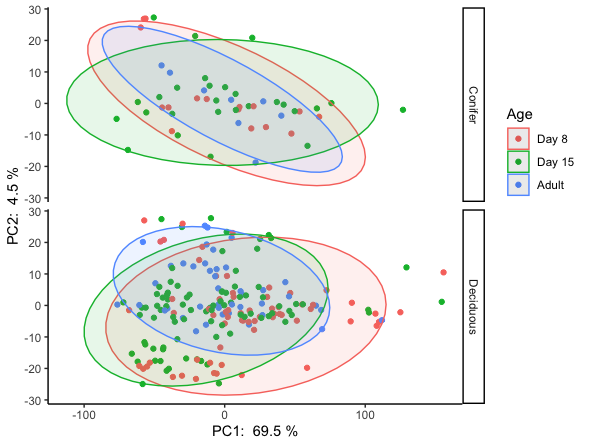
Figure S1. Scatterplot of PC1 and PC2 of microbiome community structure (beta-diversity), coloured by age and separated by habitat type. Ellipses plotted with ‘t’ distribution. Ellipses coloured by age.

Figure S2. Female Shannon diversity regressed on offspring diversity.

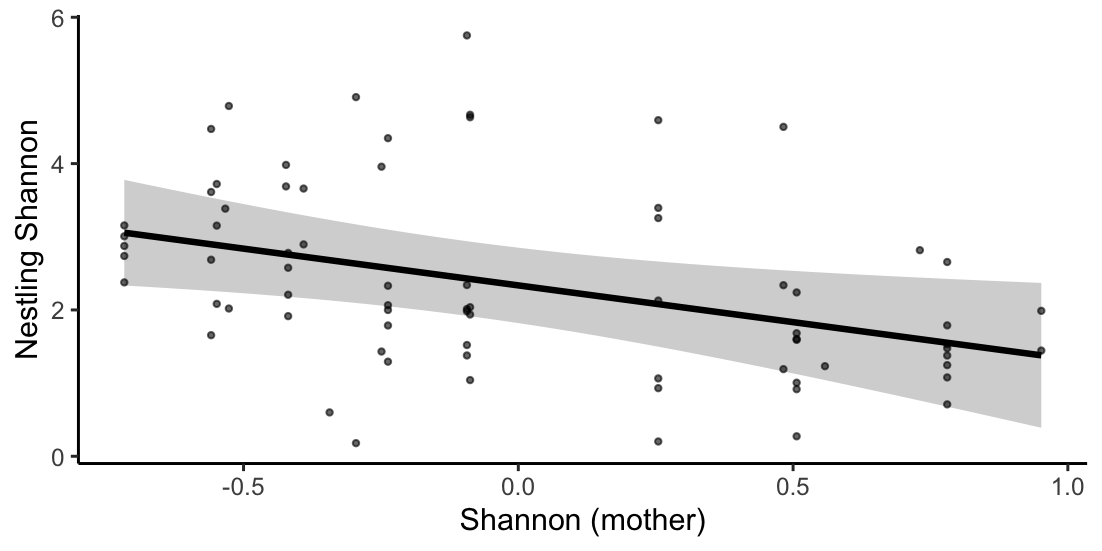
